# Isolation and Characterization of a *Halomonas* Species for Non-Axenic Growth-Associated Production of Bio-Polyesters from Sustainable Feedstocks

**DOI:** 10.1101/2024.03.28.587248

**Authors:** Sung-Geun Woo, Nils J. H. Averesch, Aaron J. Berliner, Joerg S. Deutzmann, Vince E. Pane, Sulogna Chatterjee, Craig S. Criddle

## Abstract

Biodegradable plastics are urgently needed to replace petroleum-derived polymeric materials and prevent their accumulation in the environment. To this end, we isolated and characterized a halophilic and alkaliphilic bacterium from the Great Salt Lake in Utah. The isolate was identified as a *Halomonas* species and designated “CUBES01”. Full-genome sequencing and genomic reconstruction revealed the unique genetic traits and metabolic capabilities of the strain, including the common polyhydroxyalkanoate (PHA) biosynthesis pathway. Fluorescence staining identified intracellular polyester granules that accumulated predominantly during the strain’s exponential growth, a feature rarely found among natural PHA producers. CUBES01 was found to metabolize a range of renewable carbon-feedstocks, including glucosamine and acetyl-glucosamine, as well as sucrose, glucose, fructose, and further also glycerol, propionate, and acetate. Depending on the substrate, the strain accumulated up to ∼60% of its biomass [dry w/w] in poly(3-hydroxbutyrate), while reaching a doubling time of 1.7 h at 30^◦^C and an optimum osmolarity of 1 M sodium chloride and a pH of 8.8. The physiological preferences of the strain may not only enable long-term aseptic cultivation but can also facilitate the release of intracellular products through osmolysis. Development of a minimal medium also allowed the estimation of maximum PHB production rates, which were projected to exceed 5 g_PHB_/h. Finally, also the genetic tractability of the strain was assessed in conjugation experiments: two orthogonal plasmid-vectors were stable in the heterologous host, thereby opening the possibility of genetic engineering through the introduction of foreign genes.

**IMPORTANCE:** The urgent need for renewable replacements for synthetic materials may be addressed through microbial biotechnology. To simplify the large-scale implementation of such bio-processes, robust cell factories that can utilize sustainable and widely available feedstocks are pivotal. To this end, non-axenic growth-associated production could reduce operational costs and enhance biomass productivity, thereby improving commercial competitiveness. Another major cost factor is downstream processing. Especially in the case of intracellular products, such as bio-polyesters. Simplified cell-lysis strategies could also further improve economic viability.

## INTRODUCTION

Earth’s biosphere and all lives within are being threatened by an unprecedented accumulation of synthetic materials of anthropogenic origin, commonly known as plastics [1, 2, 3]. In the US, plastics make up about 12% of municipal solid waste (MSW) [4].

Global production of petroleum-based plastics reached 391 MMt/a in 2021; projected to grow exponentially for the foreseeable future, it will soon surpass a volume of half a gigaton per year [5]. Currently, the US’ plastics industry alone accounts for 3.2 quadrillion BTU (quads) of annual energy use, resulting in over 100 Mmt CO_2_e/a of greenhouse gas (GHG) emissions [2].

Both, the contribution to global warming from the production of synthetic materials and the contamination of the biosphere at their end-of-life, have aggravated environmental problems with consequences on a global scale: the damage inflicted on the economy likely exceeds the revenue generated by the plastics manufacturing industry [6, 7]. Therefore, renewable and readily deconstructable materials are urgently needed. Hence, efforts to recycle and reuse waste streams into renewable materials are being boosted. This includes the upcycling of spent plastics and/or their synthesis from GHGs. Complementing catalytic methods, many approaches now rely upon biotechnology, employing a variety of plants, algae, fungi, or microorganism s [8, 9, 10, 11, 12, 13].

Bio-producible materials that can be obtained from renewable and waste-derived feedstocks are not only of interest to sustainable manufacturing on Earth but are also sought after for *in situ* manufacturing in space [14, 15]. For example, it has recently been estimated that about half the payload of a human deep-space exploration, which mainly consists of items related to environmental control and life-support systems, consists of plastics [16]. Enabling their production “on the go” could significantly reduce the cargo requirements of such endeavors.

Biological polyesters are promising compounds for their utilization as renewable and biodegradable thermoplastics. In the case of polyhydroxyalkanoates (PHAs), stress and/or nutrient limitation can trigger polymer synthesis from recovered carbon-feedstock [17]. Of the bio-polyesters, poly(3-hydroxybutyrate) is the most common PHA. This material has mechanical properties similar to poly(lactic acid) and can be fabricated into diverse plastic products [18, 19, 20, 21, 22, 23]. Commercial success of PHAs has, however, been hampered by their production costs, which are strongly influenced by the choice of feedstock, as well as the operational expenses (OpEx) of a biotechnological process.

Volumetric production rates often fall short of their theoretical maxima, as in most cases PHAs are only formed towards the end of the growth cycle. Further, extensive cleaning and sterilization of cultivation vessels is required between batches to avoid contamination, resulting in significant downtime. Maintaining pure cultures also requires sterile conditions throughout the run. Another cost-factor is the downstream purification of PHAs from the biomass, which commonly involves solvent-based lysis and separation steps.

Aseptic conditions can be maintained much more easily with extremophiles. In particular, halophiles have already been employed for the production of bioplastics in several instances [24, 25], as these extremophiles can be kept axenic in non-sterile environments, significantly reducing the OpEx of a process. Further, polyesters can be released from the cells through osmolysis, partially abolishing the need for solvents [26], which can save additional cost.

Here, an extremely halophilic and alkaliphilic microbe was isolated from the Great Salt Lake (GSL) in Utah and comprehensively characterized, to serve as a host for the low-cost production of polyhydroxybutyrate (PHB) from sustainable feedstocks under non-sterile conditions.

## RESULTS

### Isolation and Basic Characterization of a Halophilic Microbe

A water sample was taken from the north arm (Gunnison Bay) of the GSL (Shoshone: *Ti’tsa-pa*), as shown in FIG **1**a, from which microbes were enriched and selected for halophily under extreme osmolarity (3.4 M sodium chloride ≙ 6.8 osmol/L). Six isolates were obtained at 30^◦^C on complex substrate, all of which were belonged to the same species based on their 16S rRNA sequences (OQ359097.1). The closest relative was *Halomonas gomseomensis* M12 (NR042488.1), with ∼99.7% 16S rRNA gene identity. The hitherto undescribed *Halomonas* strain was designated “CUBES01” and exhibited a maximum growth rate of 0.402 h^−1^ on complex substrate (FIG **1**c); Nile red staining and fluorescence microscopy revealed the presence of intracellular granules (FIG **1**b), that were indicative of poly(3-hydroxybutyrate) biosynthesis.

**FIG 1.**
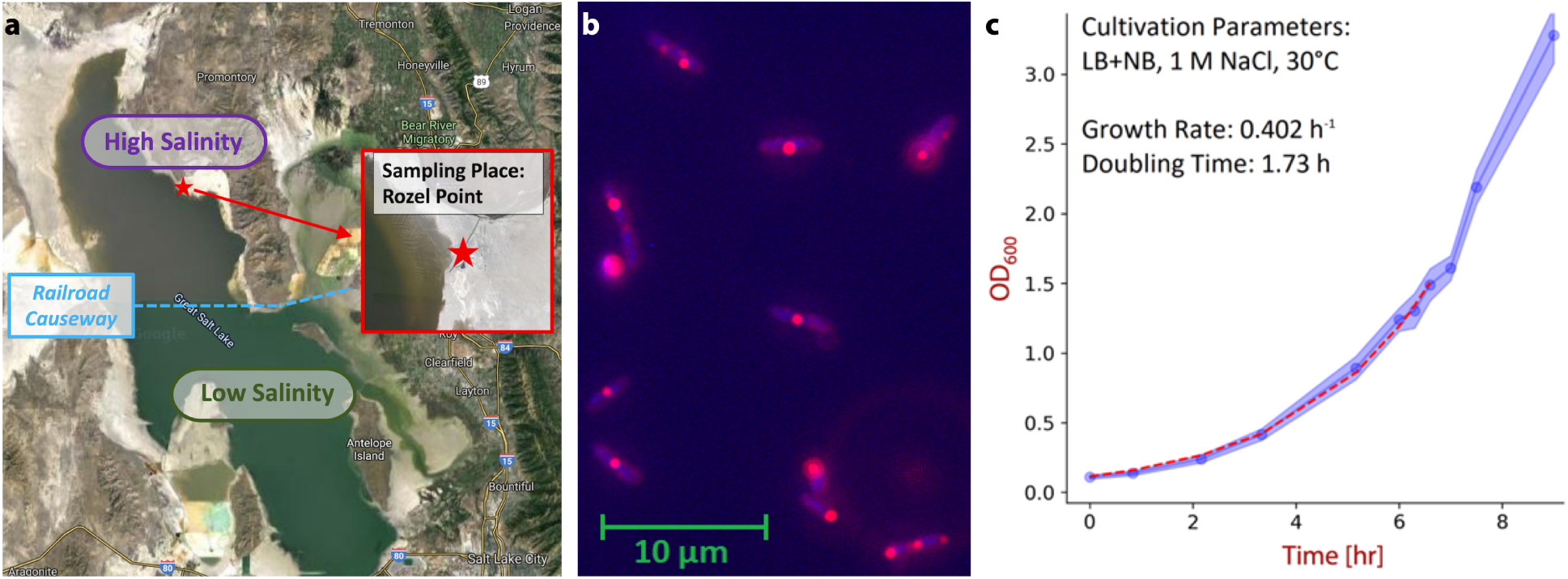
Isolation and initial characterization of *Halomonas* species. **(a)** Point of sample collection, marked on a map of the Great Salt Lake, UT, USA. Specifically, samples were taken off the Spiral Jetty near Rozel Point at coordinates 41^◦^26’15.5”N 112^◦^40’09.7”W. **(b)** Microscopy of *Halomonas* species with fluorescence-staining of intracellular polyester granules. **(c)** Growth curve of isolated *Halomonas* species on rich complex medium (aerobic, 30^◦^C, 1 M NaCl).

### Phylogenetic Classification of the *Halomonas* Isolate

The phylogenetic analysis of *Halomonas* sp. CUBES01 was conducted based on its 16S rRNA gene sequence. Within a phylogenetic tree of 50 *Halomonas* 16S RNA sequences (FIG **2**a), strain CUBES01 clustered together with *H. gomseomensis* M12 into a single branch. *H. arcis* AJ282, *H. azerica* TBZ9, *H. janggokensis* M24, and *H. subterrnea* ZG16 formed the closest neighboring group to CUBES01 and *H. gomseomensis*. The pairwise genetic distance heatmap in FIG **2**b supports the phylogenetic analysis. Specifically, *H. gomseomensis* M12 (99.7%), *H. arcis* AJ282 (98.2%), *H. subterranea* ZG16 (97.8%), and *H. janggokensis* M24 (97.7%) exhibited high 16S rRNA sequence identity. Other *Halomonas* strains exhibited a genetic identity ranging from 95.2% to 97.7%, with the exception of *H. lysinitropha* 3(2) which at 93.7% similarity was more distantly related.

**FIG 2.**
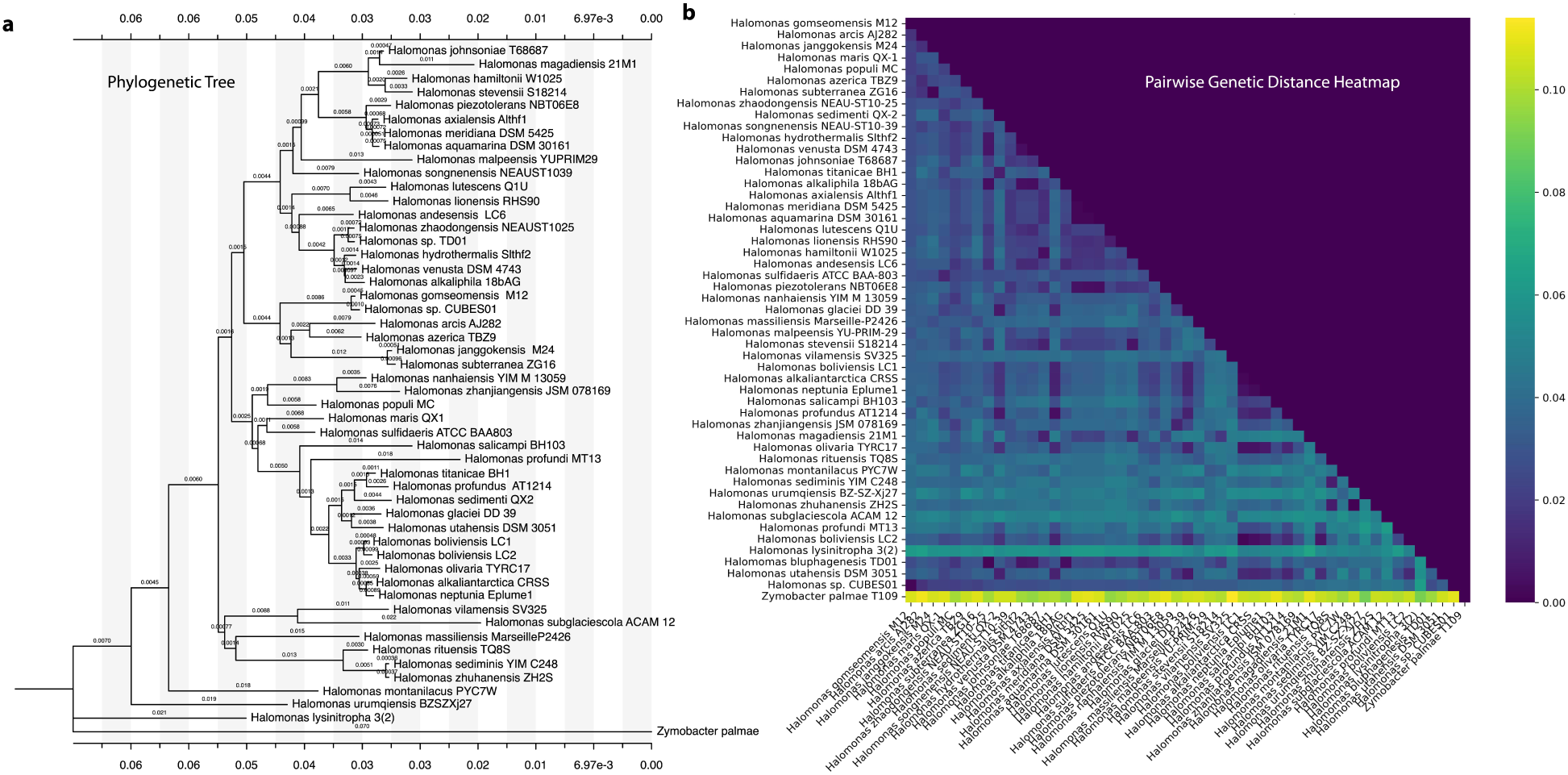
Phylogenetic affiliation and neighbourhood of *Halomonas* sp. CUBES01 based on 16S rRNA sequence comparison. **(a)** Phylogenetic Tree based on the minimum evolution (ME) method. **(b)** Heatmap of the pairwise 16S rRNA gene sequence distance of CUBES01 to other strains. Genbank IDs for each *Halomonas* species can be found in TAB S7.

The profiles of fatty acids and respiratory quinones of *Halomonas* sp. CUBES01 were consistent with the relationships observed in the pairwise genetic analysis: the predominant fatty acid, C_18:1_ *ω*7c, constituted 54.9% of the profile (see TAB S1 for full composition), the strongly predominant ubiquinone of *Halomonas* sp. CUBES01 was Co-Q9 (97.5%), followed by Co-Q8 (1.2%) and Co-Q10 (1.3%).

### Genotypic Characterization of the *Halomonas* Isolate

Full genome sequencing and genomic reconstruction of *Halomonas* sp. CUBES01 revealed a single circular chromosome of 3,642 Kbp with a G+C content of 60.1% (FIG **3**).

**FIG 3.**
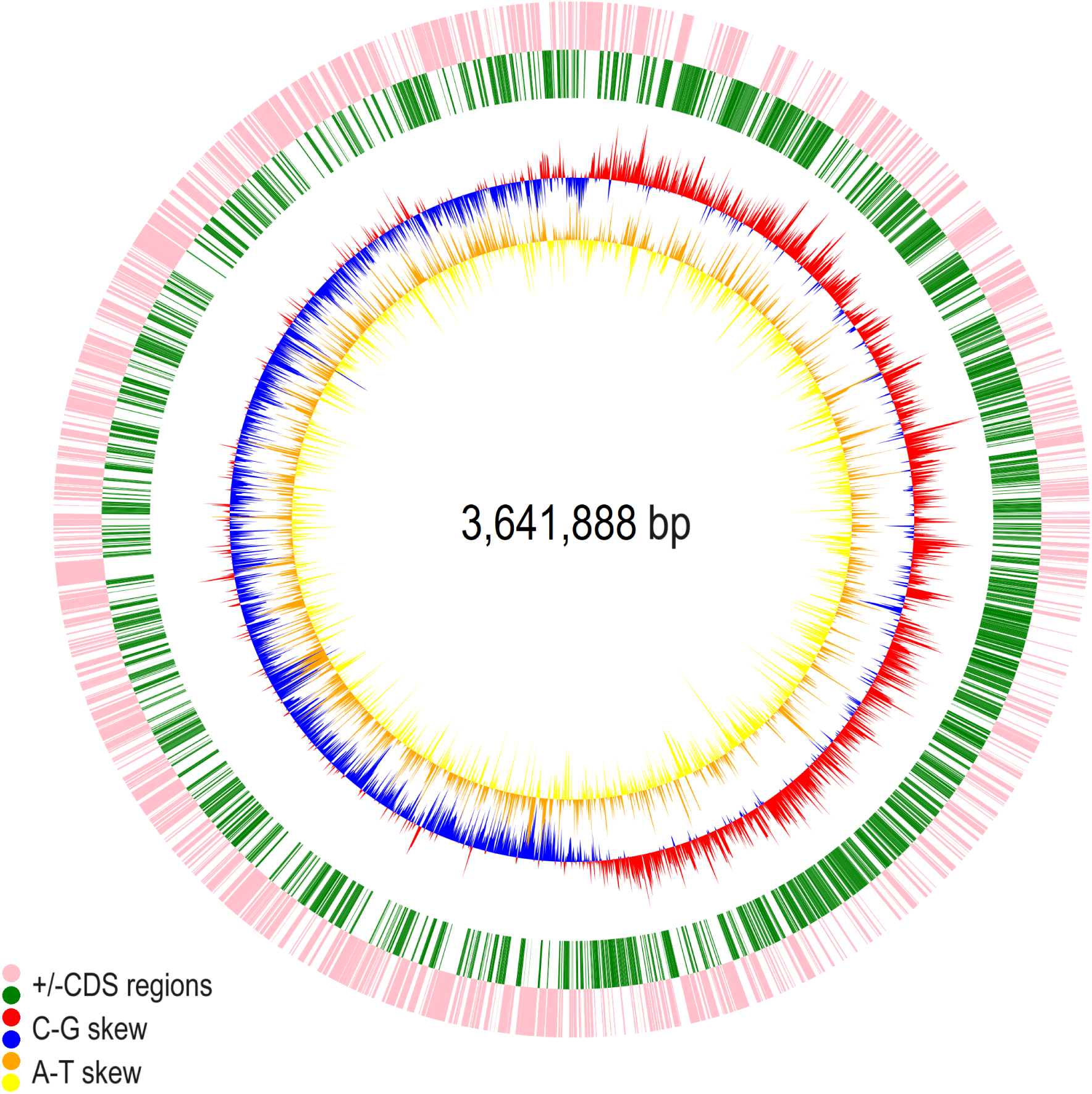
Circular representations of the *Halomonas* sp. CUBES01 chromosome displaying relevant genome features; yellow/orange: A-T skew, blue/red: C-G skew, green: +CDS regions, pink: -CDS regions.

#### Genome Features

Biological functions were assigned to 3095 of the 3396 predicted coding sequences (**TAB 1**). The remaining 301 coding sequences comprised one conserved hypothetical protein and 300 hypothetical proteins of unknown function. Preliminary analysis of the assembled genome with Pathway Tools revealed 6792 genes, 345 pathways, 2001 enzymatic reactions, 130 transport reactions, 230 protein complexes, 2193 enzymes, 684 transporters, 1369 compounds, 4786 transcription units, and 575 GO terms. In addition, the Codon Usage Bias (CUB) of *H.* sp. CUBES01 was determined based on the assembled genome files (contig 1 and 2) identified from the sequenced genome (see FIG S1).

**TABLE 1.** Features of the *Halomonas* sp. CUBES01 genome and coding sequences.

| Feature | Contig 1 | Contig 2 |
| --- | --- | --- |
| Size [bp] | 3,641,888 | 26,118 |
| Fraction of genome coding [%] | 90.41 | 92.79 |
| Coding sequences [#] | 3373 | 23 |
| Function assigned [#] | 3073 | 22 |
| Hypothetical protein [#] | 300 | 0 |
| Conserved hypothetical protein [#] | 1 | 0 |
| Average ORF length [bp] | 976.12 | 1053.65 |
| Maximal ORF length [bp] | 11895 | 2379 |
| GC content [%] | 60.00 | 60.70 |
| ATG initiation codons [%] | 87.67 | 82.60 |
| GTG initiation codons [%] | 9.07 | 8.70 |
| TTG initiation codons [%] | 3.20 | 0.0 |

#### Metabolic Features

All central metabolic pathways common among heterotrophic bacteria were annotated for *Halomonas* sp. CUBES01; a count of the major metabolic functions distinguished into 30 categories can be found in FIG S2. The genome analysis revealed the genetic basis for core catabolic capabilities, such as the uptake and breakdown of various sugars and carbonic acids and complex anabolic capabilities of potential biotechnological interest. Certain metabolic features of particular interest are highlighted in the following. A complete overview of all metabolic capabilities of *Halomonas* sp. CUBES01 is provided in the form of a metabolic map in SI2.

Species of the *Halomonas* genus are well known for their capacity to accumulate PHB [24]. In CUBES01, 17 genes associated with polyhydroxyalkanoate (PHA) synthesis were identified. Among these, three *phaA*, one *phaB*, two *phaC*, and one *phaZ* genes were annotated, encoding *β*-ketothiolase (acetyl-CoA acetyltransferase), acetoacetyl-CoA reductase, PHA synthase, and PHA depolymerase, respectively. Notably, *phaA1* (MEC4767736.1), *phaA2* (MEC4768916.1), *phaA3* (MEC4766063.1), *phaB* (MEC4766145.1), *phaC1* (MEC4767859.1), and *phaC2* (MEC4767548.1) were not organized in common operons but dispersed throughout the genome. Of the PHA synthase genes, *phaC1* belonged to class I, but *phaC2* could not be assigned to any know class (*phaC1* encoded 617 and *phaC2* 794 amino acids, corresponding to expected molecular weights of 70 kDa and 89 kDa, respectively). While the two PHA synthases are therefore not classic isozymes, both still exhibit the conserved catalytic triad (Cys-Asp-His) [27, 24]. Additionally, PHA metabolism also encompassed regulator genes such as *phaP* (MEC4767860.1) and *phaR* (MEC4766665.1) that are associated with phasin formation and PHA synthesis autoregulation, respectively, as well as one *phaZ* (MEC4768861.1) of PHA depolymerization.

The genome of CUBES01 also harbored genes of the major biosynthesis pathway of ectoine, comprised of *ectA* (MEC4767328.1), *ectB* (MEC4767329.1), *ectC* (MEC4767330.1), and *ectD* (MEC4766129.1), encoding for L-2,4-diaminobutyric acid acetyltransferase, L-2,4-diaminobutyric acid transaminase, ectoine synthase, and ectoine hydroxylase, respectively [28].

The genome of CUBES01 further contained genes associated with DABs [29] on contig 1 (MEC4768999.1 and MEC4768998.1) and contig 2 (MEC4769013.1). The DAB operon, consistent of *dabA* (PFAM: PF10070) and *dabB* (PFAM: PF00361), encodes an energy-coupled inorganic carbon (C_i_) pump) [30], presumably encoding a hydrophilic protein that is thought to establish chemical equilibrium between CO_2_ and HCO_3_^−^. Notably, the genome of CUBES01 contains one copy each of *dabA* (MEC4768999.1) and *dabB* (MEC4768998.1) genes co-located within contig 1, while another *dabA* gene (MEC4769013.1) was found within contig 2.

### Phenotypic Characterization of the *Halomonas* Isolate

Boundary conditions for cultivation of *Halomonas* sp. CUBES01 were established, enabling a more specific phenotypic characterization of the strain.

#### Basic Cultivation Conditions

*Halomonas* sp. CUBES01 exhibited growth only under conditions of substantial osmolarity: on Nutrient Broth (NB agar), biomass formed overnight at sodium chloride concentrations between 40 and 100 g/L; the lowest sodium chloride concentration enabling growth was 30 g/L (FIG S3). At 200 g/L sodium chloride, the formation of biomass only occurred after two weeks of incubation at 30^◦^C on liquid NB. Optimum growth was observed between 60 and 80 g/L sodium chloride (FIG S3), making CUBES01 a moderate halophile [31, 32]. Hence, 1 M sodium chloride was used routinely to provide optimum growth conditions (58 g/L; osmotic pressure of ∼0.3 MPa). Optimum growth was observed at 30^◦^C (growth slowed at room temperature and stagnated at 37^◦^C). The viable pH range of strain CUBES01 was determined to be between pH 7.2 and pH 9.8, with an optimum at pH 8.8 (FIG S4). CUBES01 can, therefore, also be characterized as alkaliphilic. Using the established optimum cultivation conditions (1 M sodium chloride and pH 8.8), the maximum doubling time during aerobic growth at 30^◦^C was determined as ∼1.7 h (on complex medium, FIG **1**c).

#### Substrate Spectrum

The *Halomonas* strain was further characterized by its ability to utilize different carbon sources as substrates. For this purpose, a chemically defined medium was developed (**TAB 2**). Specifically, a combination of phosphate and carbonate buffer was used to maintain alkalinity while sodium chloride upheld the required ionic strength. With potassium nitrate serving as a source of nitrogen, CUBES01 was found to accept sucrose, glucose, fructose, and glycerol, as well as acetate as carbon and energy source; in the absence of inorganic fixed nitrogen, the amino sugars glucosamine and acetyl-glucosamine served as sole carbon and nitrogen source. The highest growth rates and biomass concentrations were observed with sucrose and glucose (0.17^−1^ and 0.15^−1^, respectively). The amino sugars also enabled high cell density, albeit formed at a much slower rate (FIG **4**). The lowest cell density was obtained on fructose followed by growth on propionate. The growth rates on acetate and glycerol exceeded those on the amino sugars, but the final biomass was slightly lower. An Analytical Profile Index (API) 50 CH test further revealed the ability of the strain to utilize arabinose, galactose, maltose, mannitol, and trehalose by CUBES01 (TAB S2).

**TABLE 2.**
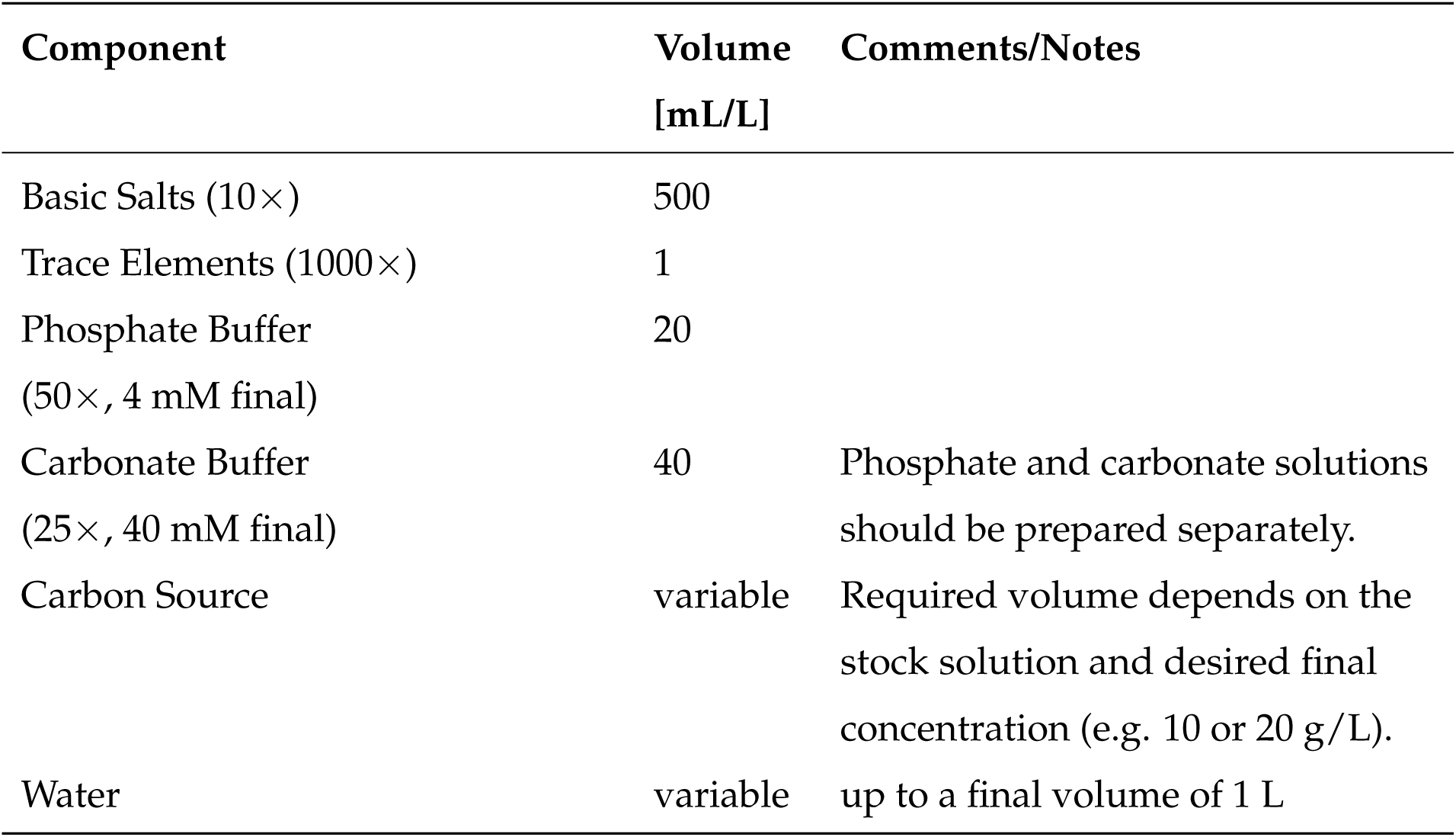
Main Components of Chemically Defined Medium.

**FIG 4.**
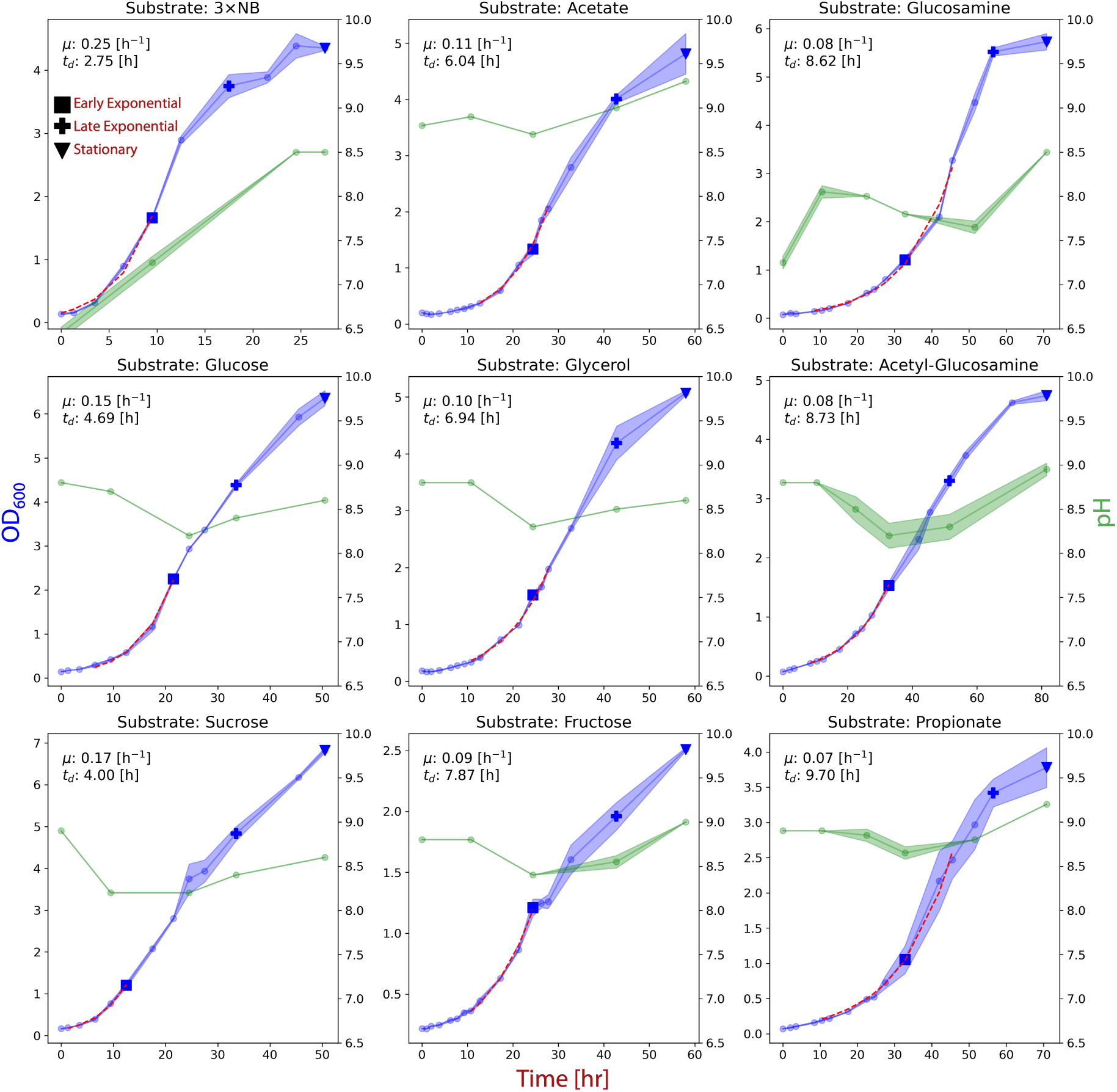
Growth of *Halomonas* sp. CUBES01 on different substrates (equivalent C-mol quantities of all substrates except for NB). The blue graphs represent the growth curves while the green graphs represent the pH throughout the cultivation. The exponential phases used to calculate growth rates are indicated by dashed red lines. Sampling points for quantification of PHB content are represented by square (early exponential), cross (late exponential), and triangle (stationary) symbols and directly correspond to the data shown in FIG **5**. The same sampling points were also used for end-product analysis.

#### Products

Using solvent extraction, polyesters can be solubilized and separated from biomass of PHA-producing microbes [33]. Applying chloroform to dried biomass of *Halomonas* sp. CUBES01, a resin was obtained that resembled a thermoplastic material when dried. Nuclear Magnetic Resonance Spectroscopy (FIG S5a) confirmed that the extracted compound was a polyester; specifically, the sextet resonance at a chemical shift of 5.25 ppm in the ^1^H-NMR spectrum was indicative of poly(3-hydroxybutyrate). Gel-Permeation Chromatography (FIG S5b) determined the weight- and number-average molecular weights (M*_w_* and M*_n_*), and polydispersity index (PDI) of the polymer, which were on the order of 572 kDa and 341 kDa, and 1.67, respectively.

Further, High-Performance Liquid Chromatography revealed the accumulation of acetate and lactate in supernatants of CUBES01 when cultivated on sugars as in the experiments underlying FIG **4**.

#### Microscopy and Morphology

While distinctively separated from each other (not adhering), live cells of *Halomonas* sp. CUBES01 appeared rod-shaped and motile, measuring approx. 1 to 4 *µ*m long and 0.8 to 1 *µ*m wide. Stained with Nile red, fluorescence microscopy revealed the count/size of intracellular inclusion bodies, which was highest/greatest during the exponential growth-phase (see FIG S6a and S6b as well as SI3 for microscopy images of samples collected at different time points during cultivation of CUBES01 on chemically defined medium as per FIG **4**).

#### Antibiotic Sensitivity and Transformability

*Halomonas* sp. CUBES01 was found to be sensitive to 43 of the 44 tested antibiotics (see TAB S3 & S4). Among these were ampicillin, carbenicillin, kanamycin, neomycin, gentamicin, tetracycline, erythromycin, streptomycin, spectinomycin, and chloramphenicol, which are commonly used as selection markers for plasmid-maintenance in shuttle-vector systems [34].

Further, to assess the strain’s genetic tractability, a protocol for the transformation of CUBES01 through bacterial conjugation was developed. Especially the strain’s strict requirement for halophilic conditions proved challenging, since common *E. coli* donor strains hardly tolerated more than 40 g/L of sodium chloride. At 35 g/L of sodium chloride, sufficient growth of both species enabled decent conjugation efficiency, yielding ∼ 100 and ∼1000 colonies per plate, respectively (depending on the vector and dilution factor, seen FIG S7).

Specifically, three different broad-host-range plasmids based on either the RP4 (RK2) or the pBBR1 conjugative system were tested (pTJS140, pBBR1MCS, and pCM66T; see TAB S5 for details). As was evident from the overnight growth of the transformed mutants, CUBES01 accepted the respective RK2/RP4, and pBBR1 plasmids pTJS140 and pBBR1MCS, selectable on Strep*^R^*/Spec*^R^* and Kan*^R^*/Neo*^R^*, well. The KanR/NeoR selectable RK2/RP4 plasmid pCM66T appeared to be difficult to maintain by CUBES01, as colony formation of transformed mutants required several days of incubation. Using PCR-based verification methods, over 90% of the isolated exconjugants of pTJS140 and pBBR1MCS could be confirmed as transconjugants (FIG S8). In addition, genetic material obtained from the cultivated *Halomonas* mutants was used to transform competent *E. coli*, verifying the presence of replicating shuttle-vector plasmids and ruling out re-arrangement, such as genomic integration of the heterologous DNA.

### Employing the *Halomonas* Isolate for Bio-Polyester Production

Isolated with the intent to be employed for sustainable bio-polyester production, the capacity of *Halomonas* sp. CUBES01 to accumulate PHB and the applicability of simplified downstream processing techniques for the release of the intracellular product were investigated.

#### Formation of PHB on Different Substrates

To further characterize the capacity of CUBES01 to form bio-polyesters, the per-biomass PHB content obtained from the cultivation of the strain on different feedstocks was assessed. Determination of the relative (normalized to biomass concentration) fluorescence intensity of cell culture stained with Nile red allowed consistent qualitative assessment of PHB production at different growth stages: on NB, accumulation of bio-polyesters appeared to be highest in early exponential phase, while on minimal medium the highest signal intensity was observed in late exponential phase for most substrates (except in case of glucose or propionate as carbon sources), as evident from FIG **5**. The highest PHB-content from gravimetric quantification after extractive purification of the bio-polyesters from biomass samples was obtained when CUBES01 was supplied with acetate, glycerol, or sucrose as substrate (69±8%, 79±8%, and 55±31% [g_PHB_/g_biomass_], respectively, as per FIG **5**).

**FIG 5.**
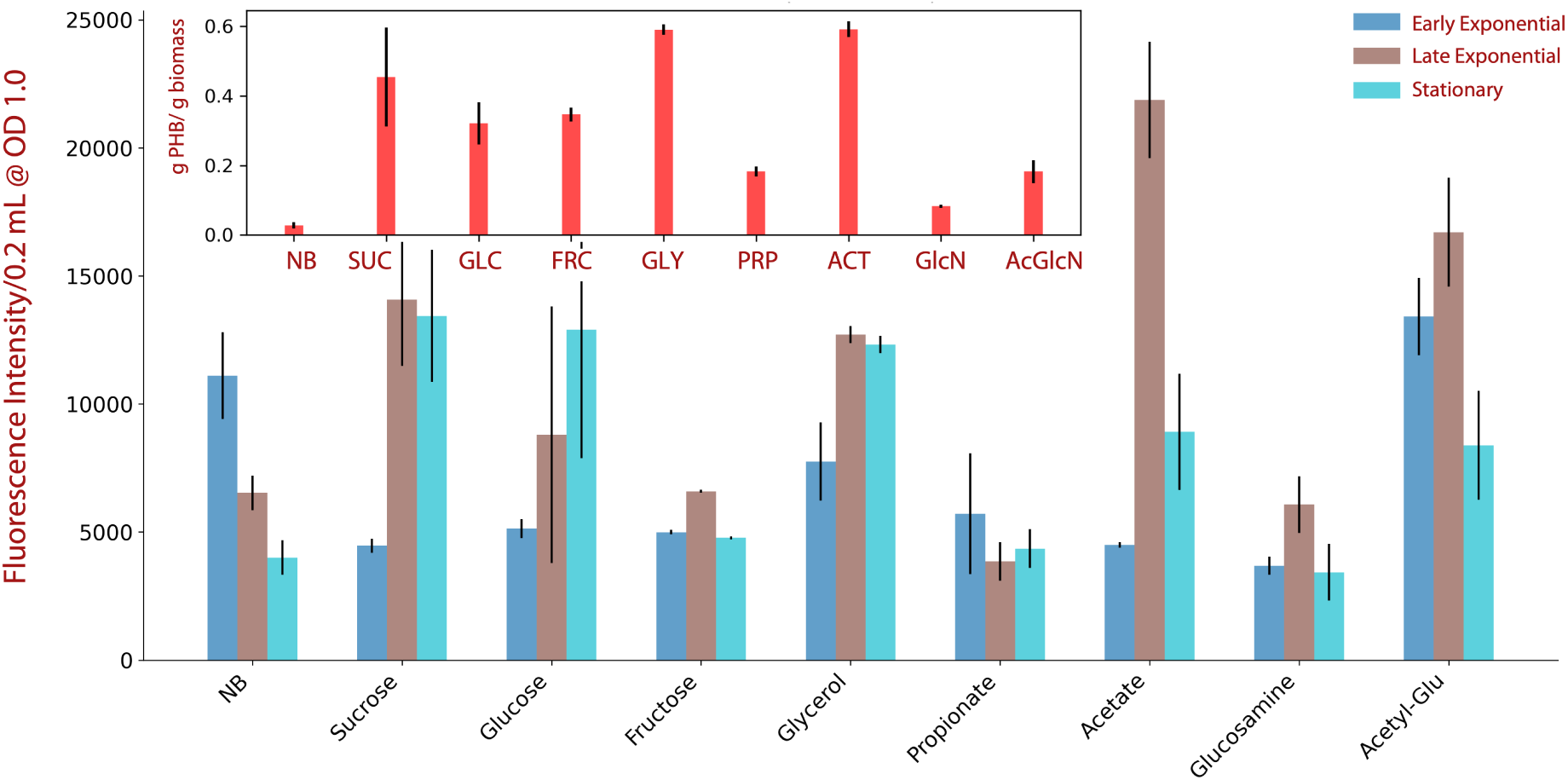
Accumulation of PHB by *Halomonas* sp. CUBES01 in early exponential (blue), late exponential (ochre), and stationary (turquoise) growth-phase on different substrates, directly corresponding to the sampling points indicated in FIG **4**. While the main chart reflects the qualitative PHB content dependent on the growth-stage (inferred by fluorescence intensity of whole-cells stained with Nile red and normalized to optical density), the insert indicates the quantitative per-biomass PHB yield (obtained gravimetrically, late exponential and stationary phase combined). NB = Nutrient Broth, SUC = Sucrose, GLC = Glucose, FRC = Fructose, GLY = Glycerol, PRP = Propionoate, ACT = Acetate, GlcN = Glucosamine, AcGlcN = Acetyl-Glucosamine.

#### Release of PHB through Osmolysis

When exposing cells of CUBES01 to deionized water, the initial optical density (OD) of 0.6 dropped almost immediately to 0.1 (FIG S9a), indicating efficient and rapid lysis. Complete lysis was confirmed by the absence of colony-forming units when dilutions of cells exposed to deionized water were plated on solid growth medium (in contrast to hundreds of colonies for the untreated culture, see FIG S9b).

## DISCUSSION

### Phylogeny

The phylogenetic tree and the pairwise distance comparisons of 16S rRNA sequences (FIG **2**) both identified *H. gomseomensis* as the closest relative of CUBES01. Analysis of the cellular fatty acid composition verified the close phylogenetic relationship between CUBES01 and *H. gomseomensis* M12 and also *H. Janggokensis* M24, which were both isolated from solar salterns in South Korea [35], and *H. lutesence* Q1T, isolated from Qinghai Lake, China [36]. Given that C16:0 was the predominant fatty acid in the profiles of *H. lysinitropha* 3(2), isolated from the Meighan Wetland, Iran, and *Halomonas* sp.

NA10-65 from the GSL, Utah [37, 38], the greater phylogenetic distance of CUBES01 from these strains is fitting. Also, the respiratory quinone pattern of CUBES01’s was consistent with those reported for *H. gomseomensis* M12, *H. janggokensis* M24, *H. lysinitropha* 3(2), and *H. lutescens* Q1U [35, 38, 36].

### Metabolism

While CUBES01 possesses all common genetic traits of PHA metabolism comprising the biosynthetic genes *phaA*, *phaB*, and *phaC*, as well as the regulatory genes *phaR* and *phaP*, they are scattered throughout the genome: the three major genes of PHA biosynthesis are not co-located in the common *phaABC* operon, only *phaP* and *phaC1* are adjacent. Common among PHA producers, this co-location often occurs within an interval of 5 to 211 nucleotides between the two genes [39]; in case of CUBES01, the distance is 112 nucleotides. In several *Halomonadaceae* the latter is often located downstream of the former (i.e. *phaP*-*phaC*, in contrast to the common organization of the PHA biosynthetic locus [40]). The coordinating role of PhaP, a so-called phasin, has been recognized previously [41]: by interacting with the N-terminal domain of PhaC, it stabilizes elongation of the polymer-chain and thus defines the size of the PHB granule. Of the two PHA synthases, PhaC1 is fairly common; PhaC2 is less common, however, homologs thereof, which have a longer C- and the shorter N-terminal domain, are still frequent in several *Halomonadaceae*, such as the species *bluephagenesis* TD01 [39]. The absence of *phaP* adjacent to the *phaC2* gene on the genome of CUBES01 in combination with the shorter N-terminal domain of that enzyme could mean that PhaP is (a) not co-expressed with PhaC2 and (b) no interaction on protein-level is possible. This could be an indication of this PHA synthase’s constitutive expression. Also PhaR, a repressor and autoregulator of PHA biosynthesis [40], was identified in CUBES01. In well-studied PHA producers, such as *Cupriavidus necator* and *Haloferax mediterranei*, PhaR is usually found close to the PHA synthetic locus and negatively regulates it, influencing PhaP expression in dependence on the abundance and maturity of PHB granules [42, 43]. In CUBES01 *phaR* is not located in the vicinity of any of the other PHA-regulatory or -biosynthetic genes. While the specific functions and interactions of PhaP and PhaR with each other as well as PHB in *Halomonas* CUBES01 remain elusive, the significant differences in genomic organization of these genes suggest a substantially different regulation of PHA metabolism. The fact that *phaA* and *phaB* are also not co-located with either of the *phaC*’s nor *phaR* and*phaP*, suggests their independent regulation, which could explain the observed perpetual PHB production.

The *dabA* genes code for a protein subunit with a domain homologous to *β*-carbonic anhydrases, which is crucial for the conversion of CO_2_ to bicarbonate (HCO_3_^−^). The *dabB*gene codes for a membrane protein that is implicated in establishing or utilizing a proton gradient, playing a role in the energy-dependent transport of inorganic carbon across the cell membrane. Together, the presence of *dabA* and *dabB* in CUBES01 distinguishes it from closely related strains commonly employed in microbial biotechnology, such as *H. bluephagenesis* TD01 [44] and *H. boliviensis* LC1 [45], as well as another isolate from the GSL north arm, *H. utahensis* DSM 3051 [46, 47]. This particular gene, implicated in inorganic carbon transport, is common among autotrophs that also bear genes such as RuBisCO and/or carbonic anhydrase [29]. Given that CUBES01’s genome does not appear to comprise a full Calvin-Benson-Bassham cycle, the presence of these genes in CUBES01 would suggest alternative roles in bicarbonate provision for essential metabolic pathways or in maintaining intracellular pH, which is connected to the metabolisms of alkaliphiles. It is also plausible that the strain lost these capabilities or that *dabA* and *dabB* were acquired through horizontal gene transfer. This logic is supported by the occurrence of two 100% identical copies of *dabA* located in both contigs.

### Phenotype & Physiology

The preference of *Halomonas* sp. CUBES01 for halophilic conditions is unsurprising, given that the isolate was first obtained from the north arm (Gunnison Bay) of the GSL where the salinity is commonly between 27 and 29%, with the predominant ions being sodium and chloride [48]. The strain’s alkaliphily, however, is unexpected, given the surveyed pH around the point of sample collection site is close to neutral [49, 50, 51].

Nevertheless, the development of a minimal (i.e., chemically defined) growth medium, derived from the optimum cultivation conditions, enabled the characterization of the strain’s growth on several carbon sources. Notably, the strain grew well on sucrose and glucose with similar biomass yield, while growing poorly on fructose, yielding significantly less biomass than on the other two sugars. This is surprising, given that sucrose is a disaccharide comprised of glucose and fructose. The reasons for this are unknown in the context of CUBES01, however, often lie with the uptake mechanisms, which in many bacteria can be rather specific to certain sugars [52, 53]. Of the non-sugar substrates, especially glycerol and acetate allowed high growth and PHB yields [54].

While both can be derived from non-edible (waste) biomass, especially the latter is seen as a next-generation feedstock in terrestrial [55] as well as space bioprocess engineering [56, 14, 15]. It is likely that the downstream entry of acetate into metabolism, which commonly proceeds via Acetyl-CoA, stimulated PHB formation, as the pathway branches off directly from that intermediate. Also, amino-sugars are of particular interest for this application, as these can be obtained from e.g., cyanobacterial lysate via *in situ* resource utilization [57]. Of these, the utilization of Acetyl-Glucosamine resulted in the highest PHB yield, which aligns with the finding that acetate enhances PHB production.

During the cultivation of CUBES01 on various substrates, we observed a consistent pattern of pH changes, characterized by an initial drop followed by a subsequent rise to near-initial levels, except in the case of complex substrate (FIG**4**). The sustained increase in pH observed with complex substrate can be attributed to its non-buffered composition, which differs from the carbonate-buffered chemically defined medium. In the latter case, CUBES01 may have initially metabolized the respective carbon sources into organic acids, such as acetate and lactate, which were found in supernatant samples. The rise of the pH during late exponential phase could be associated with a diauxic shift where the produced organic acids are consumed again [58]. This is supported by the observation that with organic acids such as acetate and propionate as substrates the final extracellular pH was slightly higher than with sugars or sugar alcohols.

### Potential of PHB Production

Interestingly, CUBES01 exhibited PHB production during both early and late growth phases, contrary to the conventional notion that PHB accumulation occurs under nutrient-depletion or electron acceptor deficient conditions [59, 24]. This underscores the multifaceted role of PHA beyond being a mere storage product for energy and carbon, extending to encompass various stress-response mechanisms. For instance, PHB production has been shown to confer salt stress resistance by preventing protein aggregation in halotolerant strains, such as *Pseudomonas* sp. CT13 [60]. Furthermore, in *Halomonadaceae*, PHB production positively correlates with salinity [61]. Another study of extreme halophiles has demonstrated the salinity threshold for halophiles’ anabolic metabolisms to exceed the presently accepted limit dictated by cell division [62]. This suggests that in halophiles PHAs may serve additional functions beyond their role as an energy storage compound. Similarly, for thermophiles such as *Chelatococcus daeguensis* TAD1, elevated heat serves as a stress factor triggering PHB accumulation during growth [63], even in the absence of nutrient limitations.

Based on the maximum growth rate and an OD-to-biomass correlation (TAB S8) that was obtained from samples collected during late exponential growth phase of CUBES01 on different substrates, biomass-specific maximum PHB production rates were estimated (FIG **6**).

**FIG 6.**
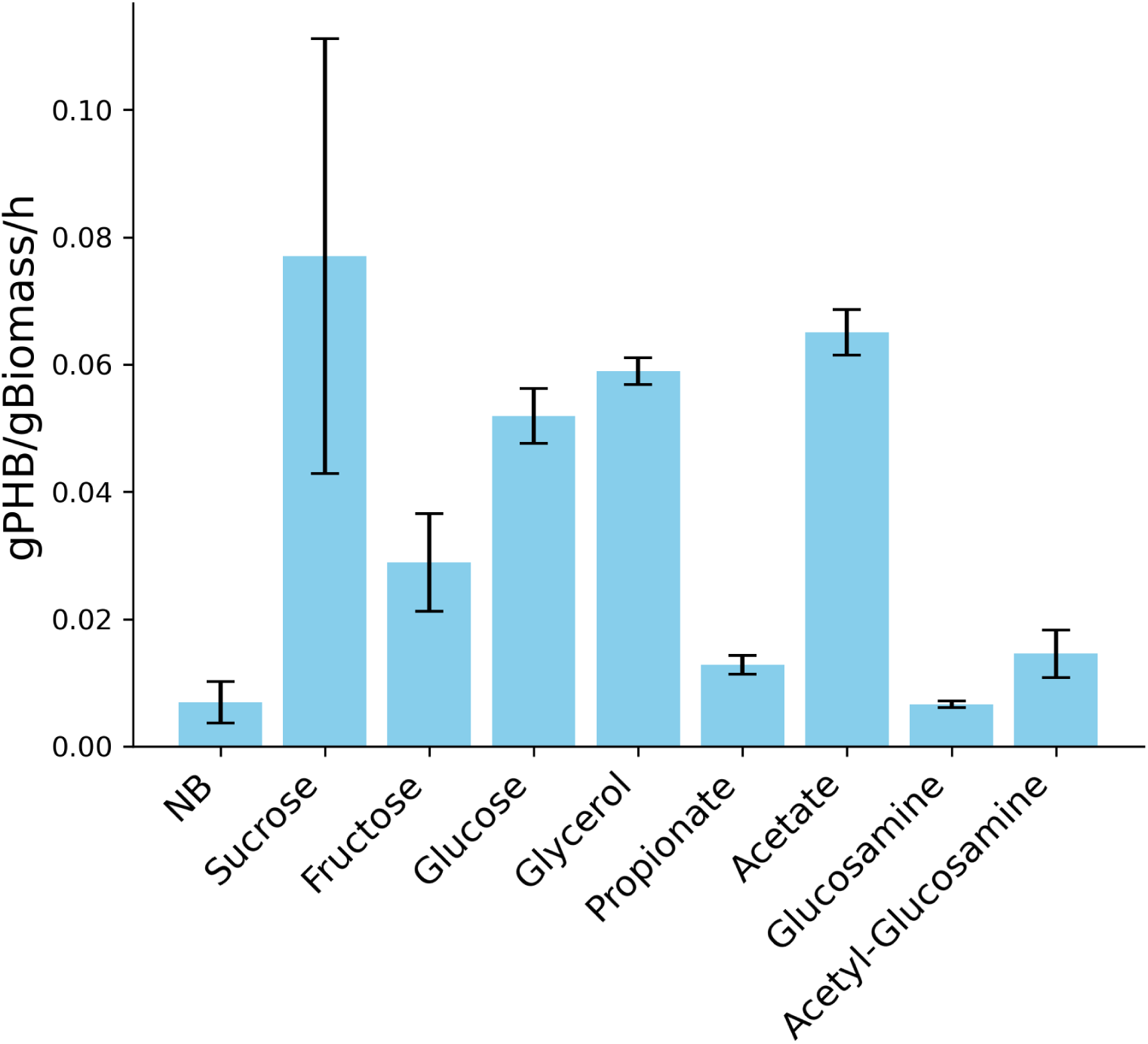
Estimated biomass-specific maximum PHB production rates of *Halomonas* sp. CUBES01. Data based on rates of biomass formation during exponential growth and PHB content during late exponential phase. Data corresponds to values shown in FIG **4** and FIG **5**. NB = Nutrient Broth, SUC = Sucrose, GLC = Glucose, FRC = Fructose, GLY = Glycerol, PRP = Propionoate, ACT = Acetate, GlcN = Glucosamine, AcGlcN = Acetyl-Glucosamine.

While the maximum overall achievable biomass concentration of CUBES01 was not specifically optimized, other *Halomonas* strains have reached maximum cell dry weights (CDW) of 87.3, 80, and 44 g/L CDW (in case of *H. venusta* KT832796, *H. bluephagenesis* TD01, and *H. boliviensis* LC1, respectively [64, 65, 66]). With an average of the reported biomass concentrations of 70.4 g/L, we deducted potential volumetric PHB production rates of 5.6, 3.5, 4.2, and 4.9 g_PHB_/h for sucrose, glucose, glycerol, and acetate respectively, based on the estimations shown in FIG **6**. These notably high projected rates suggest that CUBES01 holds great promise as a platform for the bioplastics production.

### Osmolytes and Osmolysis

Given that *Halomonas* strains thrive in hyper- or moderately saline environments, they are likely to be susceptible to lysis when subjected to rapid change osmotic downshock. Taking advantage of this phenomenon could simplify the purification of bioplastic and reduce or even entirely abolish the need for organic solvents. Mechano-sensitive genes are pivotal for imparting resistance to changes in ionic strength, as evidenced by studies demonstrating improved osmolysis when knocked out, even of non-halophilic microorganisms [67]. While such genes are also present in CUBES01 (*mscK* or *mscL*), they appear to be not preventing osmolysis susceptibility of the strain. This opens the door to simplified and rapid purification of intracellular products, reducing process complexity and resource requirements.

### Genetic Tractability

The ability to transform CUBES01 with broad-host-range plasmid-vectors through bacterial conjugation opens up the opportunity for genetic manipulation of the strain, a rare capacity among *Halomonadaceae* [68]. However, further characterization and development of genetic tools is needed before heterologous genes may be expressed or the genome of CUBES01 can be reliably modified. Poor growth of the transformed mutants bearing pCM66T was likely due to the alternate aminoglycoside 3’-phosphotransferase gene harbored by that plasmid (i.e., aph(3’)-II, while pBBR1MCS bears aph(3’)-Ia), presumably conveying insufficient antibiotic resistance. We note that aph(3’)-Ia) is also frequently annotated as *aphA1* or *kanR* and *aph*, *nptII* or *neo*, respectively.

## CONCLUSIONS

Extremophiles like *Halomonadaceae* bear great potential to make biomanufacturing more economical, as they can reduce or abolish the need for aseptic process conditions to maintain a pure culture. In addition, the high ionic strength of the cultivation medium enables the release of intracellular products, such as bio-polyesters, via osmolysis, which could be a more cost-efficient and environmentally friendly method of downstream processing. Further, the finding that CUBES01 accumulated PHB predominantly during exponential growth is significant for industrial applications, due to the possibility for continuous process operation, increasing overall productivity. Paired with projected plausible productivities in the range of 5 g/h from different carbon sources on a minimal medium, this further enhances CUBES01’s attractiveness for microbial biotechnology, improving process viability through reduced feedstock cost. The opportunity to simplify release of the intracellular product brings sustainable production of bioplastics further into reach.

## METHODS

### Sampling and Isolation of the *Halomonas* species

A 500 mL sample of water from the GSL in Utah was taken near the Spiral Jetty of Rozel Point at 41^◦^26’15.5”N 112^◦^40’09.7”W (see map in FIG. **1**a) and kept refrigerated for transport. Upon arrival at the lab 48 hours after collection, the sample was concentrated 33.3-fold by centrifuging at 4816×g for 20 minutes and re-suspension of the resulting pellet in 15 mL of the supernatant. One milliliter of the concentrated sample was used to inoculate liquid Nutrient Broth (NB) that contained 200 g/L (20% w/v) of sodium chloride. After two weeks of incubation at 30^◦^C, 100 *µ*L of the culture was spread on NB agar that contained 200 g/L of sodium chloride. The plates were incubated at 30^◦^C for two weeks. The six colonies that appeared were transferred to fresh NB agar plates with 200 g/L sodium chloride. The individual isolates were preserved as glycerol stocks for further characterization.

### Construction of Pairwise Genetic Distance Heatmap and Phylogenetic Tree

The 16S rRNA genes of the isolates were sequenced using bacterial 8F and 1492R primers as shown in TAB S6. In addition to the near full-length (1406 bp) nucleotide 16S rRNA sequence of CUBES01 (OQ359097.1), forty-nine 16S rRNA sequences of related *Halomonas* species, and one from *Zymobacter* (as an outer group), were derived from GenBank. The sequences were aligned and trimmed using Geneious by Dotmatics (Biomatters, Inc.) [69]. Based on that, the pairwise distance heatmap and the phylogenetic tree were constructed using Python 3.12.2 and MEGA11 [70], respectively. In the phylogenetic tree, the evolutionary history was inferred using the Minimum Evolution method, and the evolutionary distances were computed using the Maximum Composite Likelihood method.

### Whole-Genome Sequencing, Assembly, and Annotation

Genome sequencing of the *Halomonas* species was performed by (Plasmidsaurus, Eugene, Oregon, USA) using amplification-free long-read sequencing library preparation, utilizing Oxford Nanopore Technologies (ONT) v14 library prep chemistry. This approach was specifically chosen to ensure minimal fragmentation of the input gDNA, maintaining the integrity and continuity of the genomic sequences. A total of 570,942,477 bp were obtained across ∼148k reads. The sequencing was performed using ONT’s R10.4.1 flow cells without the use of primers. The sequencing data exhibited high quality, with the longest read being ∼66 kilobases and coverage of 155×, thus providing a robust dataset for assembly.

The genome assembly began with the removal of the bottom 5% lowest quality fastq reads via Filtlong v0.2.1 with default parameters. The reads were then down-sampled 250 megabases to create a rough sketch of the assembly with Miniasm v0.3 [71]. Using information acquired from the Miniasm assembly, the reads were then re-downsample to ∼100x coverage with heavy weight applied to removing low-quality reads. Flye [72] v2.9.1 and Medaka v1.8.0 were then leveraged to assemble with parameters selected for high-quality ONT reads. The assembled genome had a coverage of 103× encompassing 2 contigs with a combined size of 3.7 megabases. A larger contig consisted of 3,641,888 bp, while a smaller contig with higher coverage consisted of 26,118 bp, part of which was identified to be a repeat of a section of the larger contig. Functional analysis and prediction of certain genes were conducted using InterPro 98.0 [73].

Gene annotation, conducted using the Department of Energy’s KBase [74] and RASTtk v1.073 [75] using the B (Bacteria) domain default parameters, identified a feature count of 10,208 for the larger and 70 for the shorter contig. The genome analysis and characterization was based on the larger contig unless stated otherwise.

The annotated genome was imported into Pathway Tools software version 27.0 [76] where it was used to generate a Pathway/Genome Database file using the PathoLogic [77] and MetaCyc version 27.0 [78]. Pathway Tools was then leveraged to construct a comprehensive metabolic profile map (SI2).

### Analysis of Codon Usage Bias

The Codon Usage Bias (CUB) of CUBES01 was analyzed based on the assembled genome. The CUB values (frequency of codons) for each amino acid were assessed using the Jamie McGowan Bioinformatics Tools [79].

### Admittance to and Inclusion of *Halomonas* sp. CUBES01 in Strain Collections

The *Halomonas* strain was deposited to the *Deutsche Sammlung von Mikroorganismen und Zellkulturen* (DSMZ) GmbH at the Leibnitz-Institute (Braunschweig, Germany) under the designation CUBES01 and is available under accession number DSM 115203. In addition, the following services and analyses were carried out by DSMZ: Production of Biomass in Quantified Aliquots for Special Procedures, as well as Analysis of Cellular Fatty Acids (TAB S1), Analysis of Respiratory Quinones (RESULT SECTION), Analysis of Metabolic Activities (TAB S2), and Antibiotic Susceptibility Testing (TAB S3).

The *Halomonas* strain was also deposited to the Korean Collection for Type Cultures (KCTC) at the Korean Research Institute of Bioscience and Biotechnology (KRIBB) (Daejeon, South Korea) under the designation CUBES01 and is available under accession number KCTC 92801.

### Preparation of Salinity & pH Gradient-Agar

Gradient-agar plates were produced using square culture/Petri dishes, analogously to previously described toxicity tests [57]. More specifically, for salinity, two solutions of NB agar with different salt content (i.e. no sodium chloride and 175 g/L of sodium chloride, FIG. S3) were used to sequentially pour two layers of solid medium on top of each other at different angles. That way a horizontal gradient was achieved through the linear difference in vertical thickness of each individual layer. Gradient-agar plates of pH were produced similarly with layers of NB agar, as per an established protocol [80], while the total concentration of ions was kept at 1 M for optimum growth conditions. The following buffers were used to maintain the boundaries of the two different pH-gradient plates: The lower and higher pH for a gradient from 7 to 9 (FIG. S4a) was achieved with a 100 mM phosphate buffer, using the respective required amounts of KH_2_PO_4_ and KH_2_PO_4_. The lower pH for the gradient from 6.6 to 10.2 (FIG. S4b) was obtained analogously, while the higher pH was reached using a 10 mM Tris buffer system. For the latter, 90 mL of Tris solution was titrated to pH 10.2 using monovalent strong base. The volume was made up to 100 mL with pure water, achieving a final buffer concentration of 10 mM.

### Cultivation of *Halomonas* sp. CUBES01

Unless stated otherwise, the strain was routinely maintained and propagated at 30^◦^C on solid medium (agar plates) that contained 100 g/L sodium chloride with NB as the substrate. For growth experiments using liquid medium, such as to determine the maximum doubling time, as well as to accumulate biomass for PHB production, *Halomonas* sp. CUBES01 was cultivated at 30^◦^C in 500 mL baffled shake-flasks (polycarbonate with vented screw-cap) with 180 rpm shaking (culture volume maximum 10% of shake-flask capacity). Unless stated otherwise, 1 M sodium chloride maintained optimum osmotic strength, e.g. when using Luria-Bertani (LB), Nutrient Broth (NB), or a 1:1 mixture of both as substrate. The composition of the chemically defined (minimal) medium used to characterize the growth of *Halomonas* sp. CUBES01 on different substrates and to determine the biomass- and PHB yields, as well as for analysis of extracellular metabolites, is given in **TAB 2** while the composition of the respective stock solutions is provided in **TAB 3**.

**TABLE 3.** Composition of Stock Solutions for Chemically Defined Medium.

| Component | Amount [g/L] | Comments/Notes |
| --- | --- | --- |
| Basic Salts |  |  |
| KNO<sub>3</sub> | 2 | Prepare as 2× stock solution and autoclave at 121°C for 20 minutes. Let cool down before adding remaining components. |
| MgSO<sub>4</sub>·7H<sub>2</sub>O | 0.4 |  |
| CaCl<sub>2</sub>·2H<sub>2</sub>O | 0.04 |  |
| NaCl | 116.88 |  |
| Trace Elements |  |  |
| Na<sub>2</sub>EDTA | 5 | Prepare as 1000× stock solution and autoclave at 121°C for 20 minutes. After autoclaving the solution becomes pink. |
| FeSO<sub>4</sub>·7H<sub>2</sub>O | 2 |  |
| ZnSO<sub>4</sub>·7H<sub>2</sub>O | 0.3 |  |
| MnCl<sub>2</sub>·4H<sub>2</sub>O | 0.03 |  |
| CoCl<sub>2</sub>·6H<sub>2</sub>O | 0.2 |  |
| CuSO<sub>4</sub>·5H<sub>2</sub>O | 1.2 |  |
| Na<sub>2</sub>O<sub>4</sub>W·2H<sub>2</sub>O | 0.3 |  |
| NiCl<sub>2</sub>·6H<sub>2</sub>O | 0.05 |  |
| Na<sub>2</sub>MoO<sub>4</sub>·2H<sub>2</sub>O | 0.05 |  |
| H<sub>3</sub>BO<sub>3</sub> | 0.03 |  |
| Phosphate Buffer (80 mM) |  |  |
| KH<sub>2</sub>PO<sub>4</sub> | 5.44 | Prepare as 50× stock solution (total phosphate concentration of 80 mM) and autoclave at 121°C for 20 minutes, the pH of the solution should be between 6.8 and 7. |
| Na<sub>2</sub>HPO<sub>4</sub> | 5.68 |  |
| Carbonate Buffer (1 M) |  |  |
| NaHCO<sub>3</sub> | 75.6 | Prepare as 25× stock solution (total carbonate concentration of 1 M) and sterilize by filtration only, the pH of the solution should be between 8.6 and 9. |
| Na<sub>2</sub>CO<sub>3</sub> | 10.5 |  |
| Carbon Source |  |  |
| Variable (Sucrose, Glucose, Fructose, Glycerol, Glucosamine, Acetate, Acetyl-Glucosamine) |  | Prepare suitable stock solutions (e.g. 10× or 20×) for desired concentration (e.g. 100 or 200 g/L). Sterilize by filtration only. |

Microbial growth in liquid culture was monitored by determining the optical density at a wavelength of 600 nm (OD600) using a DR2800™Portable Spectrophotometer (HACH). Culture samples were collected by centrifugation (4480×g for 20 minutes) to obtain supernatant for metabolite analysis and biomass for PHA extraction. Generally, experiments involving shake-flask cultivation were performed in duplicates.

### PHA Extraction and ^1^H Nuclear Magnetic Resonance (NMR) Spectroscopy

Bio-polyesters (PHAs) were recovered from freeze-dried cell-mass employing chloroform extraction [33]. The weight of the recovered polymer was determined gravimetrically and the composition was analyzed employing Nuclear Magnetic Resonance (NMR) spectroscopy as previously reported [81]: A few mg of polymer were dissolved in deuterated chloroform and ^1^H-NMR spectra were recorded at 25^◦^C on a Unity INOVA™500 NMR Spectrometer (Varian Medical Systems) with chemical shifts referenced in ppm relative to tetramethylsilane.

### Gel-Permeation Chromatography (GPC)

Gel-Permeation Chromatography (GPC) was carried out in chloroform on a TSKgel SuperHZM-H column (Tosoh) with a DAWN MultiAngle Light Scattering (MALS) detector (Wyatt Technology) and an Optilab T-rEX differential refractometer (Wyatt Technology). Polystyrene calibrated (from *M_p_* = 500 − 275, 000 g/mol) molecular weights were determined using a GPCmax autosampler at 25^◦^C at a flow rate of 1 mL/min.

### High-Performance Liquid Chromatography (HPLC)

Quantification of acetate and lactate in the culture-broth was based on a previously published HPLC method for the detection of organic acids [82]. Briefly, the procedure was as follows: Samples of 1 mL were filtered (PVDF or PES syringe filters, 0.2 *µ*m pore-size) and diluted 1:100 into HPLC sampling vials. Analysis of 50 *µ*L sample volume was performed on a 1260 Infinity HPLC system (Agilent), using an Aminex HPX87H column (BioRad) with 5 mM H_2_SO_4_ as the eluent, at a flow rate of 0.7 mL/min. Organic acids were identified by their retention times and quantified by comparison to standards of known concentration using a refractive index detector operated at 35^◦^C or a UV detector at 210 nm.

### Microscopy and Fluorescence Staining of Intracellular PHAs

Cells previously frozen in 20% glycerol were thawed on ice, and 10 *µ*L were stained with 2 *µ*L of a 10 *µ*g/mL Nile-Red solution. Microscopy was performed on a Leica DM 4000 B epifluorescence microscope using an HCX PL APO 100x oil immersion objective with identical illumination and magnification settings for all images. Images were taken with a Leica DFC 500 camera and the Leica Application Suite V 3.8 software with identical settings for all images. Scale bars were added manually based on the camera software’s information that one pixel represents 0.092 *µ*m.

For determination of the relative fluorescence intensity of the cell-cultures, a Spark®Multimode Microplate Reader (TECAN) was used. Samples previously collected from different growth-stages were diluted to an approx. OD600 of 1 as applicable and aliquots of 200 *µ*L containing 0.0001% Nile red were distributed into a clear 96-well round-bottom microtiter plate, along with no-growth (media) blanks. The optical density and fluorescence were measured in four runs, using the following specific settings: Absorbance at 600 nm wavelength with 10 flashes and 50 ms settle time across all runs. Fluorescence by monochromator (excitation and emission) across all runs;

- bottom reading: 30 flashes (5×6 flashes per well), 40 *µ*s integration time, 0 *µ*s lag time, 0 ms settle time, Z-position of 26 mm with either:

**–** 530 nm excitation wavelength, 20 nm excitation bandwidth, 610 nm emission wavelength, 20 nm emission bandwidth, gain optimal of 109;
**–** 535 nm excitation wavelength, 10 nm excitation bandwidth, 610 nm emission wavelength, 20 nm emission bandwidth, gain optimal of 126;
- top reading: 30 flashes, automatic mirror (dichroic 560), 40 *µ*s integration time, 0 *µ*s lag time, 0 ms settle time, Z-position of 20 mm with either:

**–** 530 nm excitation wavelength, 20 nm excitation bandwidth, 610 nm emission wavelength, 20 nm emission bandwidth, gain optimal of 55;
**–** 535 nm excitation wavelength, 10 nm excitation bandwidth, 610 nm emission wavelength, 20 nm emission bandwidth, gain optimal of 80;

The individual reads were calibrated to zero based on the recorded baseline absorbance of the respective blanks for each respective medium and run, and normalized to the absorbance before averaging the reads of the samples of the four runs; the standard deviation of the samples was calculated for the biological replicates of the shake-flask cultures.

### Osmolysis Susceptibility Test

The lysis-rate of CUBES01 when exposed to an osmotic shock was quantified as the change of optical density (OD600) of a cell-suspension over a short time. Cells from the exponential phase of shake-flask culture on NB-saline medium (1 M of sodium chloride) were harvested by centrifugation (4480×g for 20 minutes) and resuspended in deionized water, while the OD was measured before the osmotic shock, as well as at 1 min, 10 min, 20 min, and 40 min thereafter. Further, colony-forming units (CFU) of the original culture, as well as the cell-suspension after the osmotic shock were determined by plating 100 *µ*L of in 10^4^-fold dilutions, based on the initial OD600 (0.6) of the culture at the point of collection, which had an estimated cell-count of ∼ 4.7×10^7^ cells/mL [83].

### Transformation of CUBES01 by Conjugation

The protocol for transformation of *Halomonas* sp. CUBES01 was developed outgoing from an established method for plasmid-vector mobilization by means of bacterial conjugation, with certain conditions adjusted in analogy to protocols for other halophiles [84], as outlined in the following. Specifically, the B2155 derivative *E. coli* strain WM3064 (*thrB1004 pro thi rpsL hsdS lacZ*ΔM15 RP4-1360 Δ(*araBAD*)567 Δ*dapA*1341::[*erm*-*pir*]), an RP4 mobilizing *λpir* cell-line which is auxotrophic for diaminopimelic acid (DAP), was transformed with the respective plasmid vectors (pTJS140, pBBR1MCS, and pCM66T; see TAB. S5 for details) using the “Mix and Go!” *E. coli* Transformation Kit (Zymo Research, Irvine, CA).

The WM3064 strains bearing the respective plasmid-vectors, hereafter referred to as the donor strains, were incubated at 30^◦^C overnight on solid LB containing DAP (300 *µ*M) and kanamycin (50 *µ*g/mL) or streptomycin (50 *µ*g/mL) as applicable, while the recipient strain, CUBES01, was incubated at 30^◦^C overnight on solid NB with 1 M sodium chloride. From the agar-plates, liquid cultures of the donor strains were inoculated on LB containing the respective antibiotic and DAP, incubated with shaking at 30^◦^C overnight. Simultaneously, the recipient strain was inoculated in the liquid cultures of NB with 1 M sodium chloride, also incubated at 30^◦^C overnight. On the next day, 6 *µ*L of the donor-strain culture was used to inoculate 3 mL of fresh LB containing DAP and 35 g/L of sodium chloride (no antibiotics). Likewise, 20 *µ*L of the recipient strain was used to inoculate 10 mL of fresh NB containing 35 g/L of sodium chloride. Both cultures were incubated with shaking at 30^◦^C. After four hours the two liquid cultures (3 mL of the donor strain and 10 mL of the recipient strain) were combined, and the cells were collected by centrifugation (10 min at 4816×g). The supernatant was discarded, and the cells were resuspended in the remaining liquid. The cell-suspension was pipetted as a drop on a prewarmed NB plate containing DAP and 35 g/L sodium chloride and incubated at 30^◦^C overnight with the plate facing up. On the next day, the biomass was collected and suspended in 500 *µ*L of NB that contained 1 M of sodium chloride. Aliquots of the cell-suspension were diluted 1:10 and 1:100, and volumes of 100 to 200 *µ*L of all three concentrations (original suspension and two dilutions) were plated on NB that contained 1 M sodium chloride and the appropriate antibiotic for the respective plasmid-vector. The plates were incubated at 30^◦^C and single colonies that appeared after 1-2 days were isolated on the same type of solid medium for screening purposes.

### Validation of Transformed CUBES01

DNA was purified from the antibiotic-resistant mutants of *Halomonas* sp. CUBES01 using a Plasmid Mini Kit (QIAGEN, Hilden, Germany) for screening on the presence of the respective plasmids. The amplicons indicative of the shuttle vectors were obtained from PCRs targeting complementary regions of the respective plasmids: two primer-sets were used per plasmid; (i) araC (forward) and spc (reverse), (ii) spc (forward) and araC (reverse) were used to confirm the presence of pTJS140; (i) neoR (forward) and araC (reverse), (ii) araC (forward) and neoR (reverse), were used to confirm the presence of pBBR1MCS.

## DATA & CODE AVAILABILITY

The sequenced genome of *Halomonas* sp. CUBES01 was deposited to the GenBank database under accession number ASM2099100v2. The GenBank accession number for the 16S rRNA gene sequence of CUBES01 is OQ359097.1. All other data and code are freely accessible as a GitHub repository.

## ACKNOWLEDGEMENTS

This work was partially supported by the National Aeronautics and Space Administration (NASA) under grant or cooperative agreement award numbers NNX17AJ31G and 80NSSC22K1474. Any opinions, findings, conclusions, or recommendations expressed in this material are those of the author and do not necessarily reflect the views of NASA.

